# Optimising ddRAD sequencing for population genomic studies with ddgRADer

**DOI:** 10.1101/2022.10.08.508655

**Authors:** Aparna Lajmi, Felix Glinka, Eyal Privman

## Abstract

Double-digest Restriction-site Associated DNA sequencing (ddRADseq) is widely used to generate genomic data for non-model organisms in evolutionary and ecological studies. Along with affordable paired-end sequencing, this method makes population genomic analyses more accessible. However, multiple factors should be considered when designing a ddRADseq experiment, which can be challenging for new users. The generated data often suffer from substantial read overlaps and adaptor contamination, severely reducing sequencing efficiency and at times affecting data quality. Here, we analyse diverse datasets from the literature and carry out controlled experiments to understand the effects of enzyme choice and size selection on sequencing efficiency. The empirical data reveal that size selection is imprecise and has limited efficacy. In certain scenarios, a substantial proportion of short fragments pass below the lower size-selection cut-off resulting in low sequencing efficiency. However, enzyme choice can considerably mitigate inadvertent inclusion of these shorter fragments. A simple model based on these experiments is implemented to predict the number of genomic fragments generated after digestion and size selection, number of SNPs genotyped, number of samples that can be multiplexed, and the expected sequencing efficiency. We developed ddgRADer - http://ddgrader.haifa.ac.il/ - a user-friendly webtool that aids ddRADseq experimental design while optimising sequencing efficiency. This tool can also be used for single enzyme protocols such as Genotyping-by-Sequencing (GBS). Given user-defined study goals, ddgRADer recommends enzyme pairs and allows users to compare and choose enzymes and size-selection criteria. ddgRADer improves the accessibility and ease of designing ddRADseq experiments and increases the probability of success of the first population genomic study conducted in labs with no prior experience in genomics.

## Introduction

Restriction-site Associated DNA sequencing (RADseq; Baird et al., 2008) is a widely used method to generate reduced representation genomic libraries. This approach dramatically reduces the genotyping cost per sample compared to whole genome sequencing, thereby allowing a limited budget project to achieve larger sample sizes, which are the most important factor determining statistical power of population genomic analyses. RADseq has been particularly beneficial in SNP discovery and genotyping of non-model organisms for population, ecological, evolutionary, and conservation genomic studies (Clugston et al., 2019; Davey et al., 2011; Eaton & Ree, 2013; Luikart et al., 2003; Wagner et al., 2013). The key advantage of this method is the flexibility it offers in the number of cut-sites that can be targeted across the genome, through the choice of appropriate restriction enzymes. This allows for a wide range of applications, from demographic surveys and population structure analyses that guide conservation efforts (for example Natesh et al., 2017; Zecherle et al., 2021), to high-resolution scans looking for the genomic basis of local adaptation (for example Magalhaes et al., 2021).

Numerous methods of generating reduced representation libraries are available that offer different advantages and help deal with different biases in data (reviewed in Andrews et al., 2016). Double-digest RAD sequencing (ddRADseq) is one of the most commonly used variations of RADseq, where genomic DNA is digested with two different restriction enzymes, generating genomic fragments in a reproducible manner (Peterson et al., 2012). Typically, a combination of a rare- and a frequent-cutting enzyme is used. The rare cutter determines the number of fragments sequenced and the frequent cutter determines the average length of these fragments. While the general rule of thumb is that enzymes that recognise shorter target sequences cut more often, the GC content in the recognition sequence also affects the number of cut sites. When choosing enzymes for a ddRADseq experiment, the size and GC content of the genome, and the polymorphism expected in the study population should be taken into consideration as they determine the number of polymorphic loci that will be genotyped, the depth per locus, and hence the number of samples that can be multiplexed in a single lane of sequencing. Taking these numerous factors into account can make designing a ddRADseq study challenging for beginners and easily result in a suboptimal study design even for experienced researchers. This can lead to substantial waste of sequencing effort. Moreover, it can generate sequencing data that is ill-fit for the downstream analyses and study goals (Puritz et al., 2014). Given the wide applicability of this method, it is crucial to make designing ddRADseq experiments more accessible and aid researchers in optimal experimental design.

Two of the biggest challenges in using ddRADseq are lack of appropriate training in designing experiments and the overall high cost of the project (compared to Sanger sequencing or microsatellites genotyping), despite the relatively low per-sample cost (compared to whole genome sequencing). Given the large capacity of a single lane of Illumina sequencing, most projects tend to conduct only one sequencing run on all samples together, multiplexed in one or a few sequencing lanes. Thus, unlike Sanger sequencing experiments, most researchers cannot afford sequential improvements to the experimental design. These challenges are particularly consequential for researchers in developing regions, including the tropics where most of the biodiversity lies and conservation efforts are needed. Several tools are available for processing of ddRADseq data and SNP calling (for example J. Catchen et al., 2013; J. M. Catchen et al., 2011; Eaton & Overcast, 2020; Nadukkalam Ravindran et al., 2019; Puritz et al., 2014; Sovic et al., 2015). However, these applications are meaningful only when the sequence data are generated with an appropriate experimental design. Relatively fewer tools can help in designing and optimising the experimental design (but see Lepais & Weir, 2014; Mora-Márquez et al., 2017; RiveraLColón et al., 2021). While these tools may help in enzyme choice and size-selection, they are based only on predictions generated in silico that may be unrealistic.

With the advent of Illumina paired-end sequencing, where reads are sequenced from both ends of a DNA fragment, twice the amount of data can be generated at a relatively low additional cost compared to single-end sequencing. Thus, paired-end sequencing is nowadays the common practice. When carrying out paired-end sequencing of 150bp reads, if the DNA fragment (insert size) is less than 300bp (two read lengths), the ends of the reads overlap, and some genomic positions are sequenced twice. If the insert size is less than 150bp (one read length), the adaptors are inadvertently sequenced leading to adaptor contamination at the end of each read, in addition to complete overlap. Any sequenced fragments shorter than two read lengths lead to considerable loss of sequencing effort apart from a dataset contaminated with adaptors. RADseq protocols include measures to remove short fragments using magnetic beads and gel-based size selection (e.g., the Bluepippin instrument). The magnetic beads selectively bind to DNA fragments typically between 200 to 500bp, but they do not completely exclude short fragments. It is possible to alter the bead-to-sample ratio to shift the minimum size threshold up or down, but this flexibility is rather limited compared to gel-based size selection. When using only magnetic beads for size selection, a previous study in our lab (Inbar et al., in prep) resulted in 74% of the sequenced read pairs having overlaps (i.e., fragments shorter than two read lengths). As a result, 27% of sequenced bases were wasted, that is, sequencing efficiency of only 73%. While the data was adequate to address the question in hand, this massive waste of sequencing effort motivated us to conduct a methodological study to optimise our sequencing efforts.

In this paper, we describe a user-friendly webtool that we developed based on our methodological insights, to assist researchers in choosing enzyme pairs and size-selection criteria while minimising waste due to short fragments. The tool allows researchers to digest their genome of interest in silico, compare enzyme combinations and size-selection cut-offs, calculate the number of samples to be multiplexed and predict the extent of adaptor contamination and read overlaps. The development of this tool was based on our work to optimise ddRADseq experimental design by minimising adaptor contamination and overlapping reads. We conducted a literature survey to look at the common choices of enzyme combinations, the size-selection methods and cut-offs used in ddRADseq studies. We re-analysed 12 of these datasets to understand the relationship between empirical adaptor contamination and read overlaps, and the theoretical expectations from in silico digestion. We conducted controlled experiments to test switching the frequent cutting enzyme to an enzyme with less cut sites, to increase the fragment sizes and hence reduce the waste and improve sequencing efficiency. This led to insights into the efficacy of size selection, allowing more realistic predictions for the implications of alternative enzyme choices and size selection cut-offs. We hope that the insights from this study and their implementation in a user-friendly webtool will aid inexperienced researchers to maximise their ddRADseq sequencing efficiency and optimise their experimental design.

### Webtool: ddgRADer

ddgRADer (http://ddgrader.haifa.ac.il/) was built to optimise sequencing efficiency by aiding enzyme and size-selection choices. It was implemented as a webtool to make designing ddRADseq experiments more accessible. This tool can also be used for other RADseq protocols that involve one or two enzymes and no random shearing such as Genotyping-by-Sequencing (GBS; Elshire et al., 2011) or ezRAD (Toonen et al., 2013).

DdgRADer works on a user-provided reference genome along with some additional basic information. The genome is in silico digested (python scripts “HandleFastafile.py” and “DigestSequence.py”) with either a predetermined list of 14 commonly used enzyme pairs or enzyme combinations chosen by the user from a comprehensive list (see the two usage modes “I have no idea!” and “Lemme try out!” in Fig. 1). Based on the nature of the question being addressed, the desired number of SNPs to be genotyped (for example to carry out demographic analysis) or the desired SNP density (for a genomic scan analysis) are specified by the user. The user also enters an estimate of the expected SNP density in the study population, which is needed to predict the number of SNPs that will be genotyped by ddRADseq (the user guide provides some examples for reference). Additional input information includes read length, single- or paired-end sequencing, expected yield from one sequencing lane, and the desired average sequencing depth for each sample. Details of the ddgRADer code are provided in the README file at https://github.com/felixglinka/ddRadSeqWebTool/blob/development/README.md

**Figure 1:**
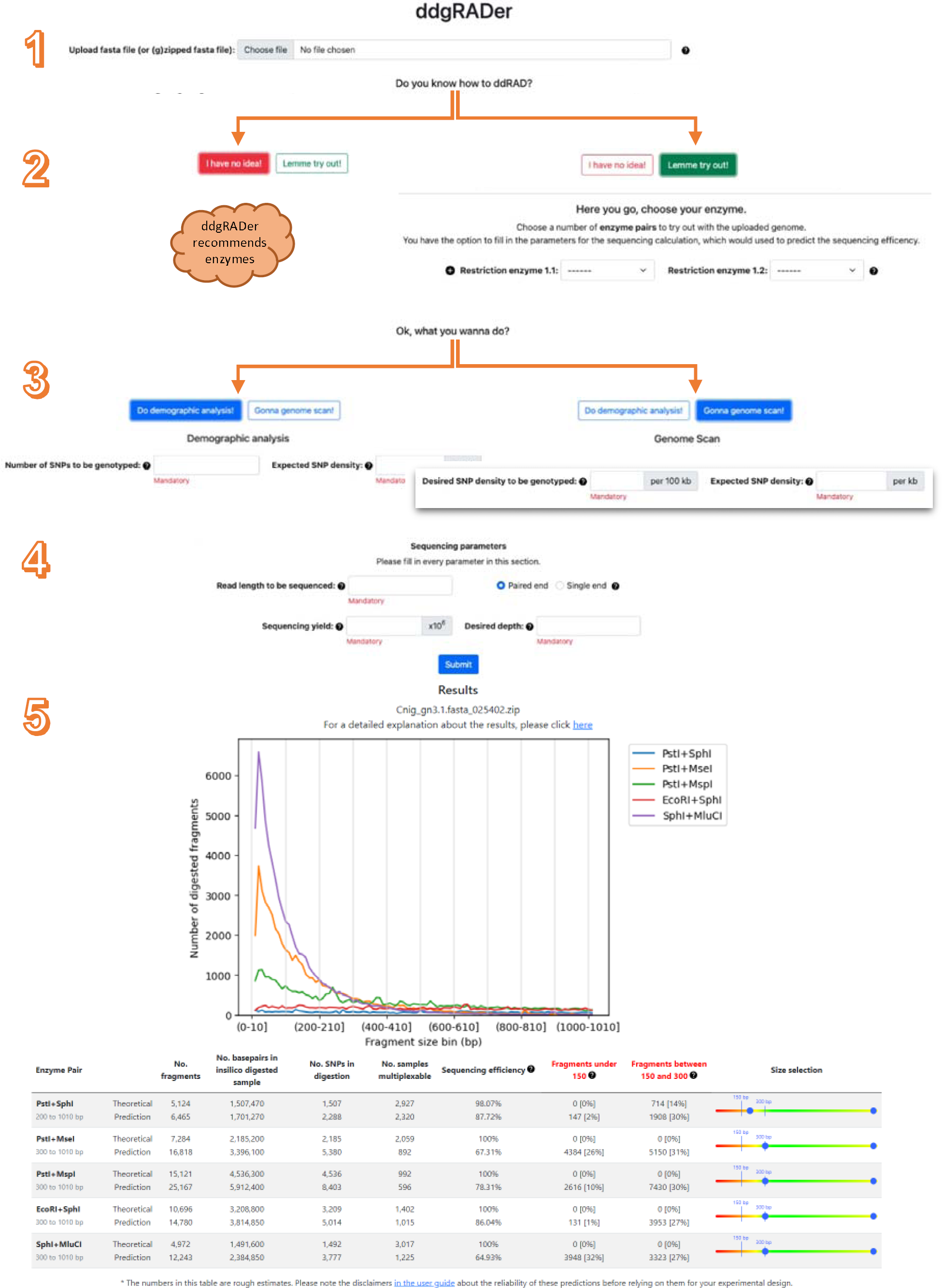
Overview of the workflow of the ddgRADer webtool and output of ddgRADer for the genome of Cataglyphis niger with five different enzyme combinations (with 150bp paired-end sequencing, 300M expected yield, and 20X desired depth). The user uploads the reference genome sequence and enters numerical parameters including the desired number of SNPs and sequencing parameters. The output consists of the in silico predicted fragment size distribution and a table that lists theoretical (based on in silico digestion) and predicted (based on the empirical studies) values from the webtool.

The in silico digestion results in the expected number of fragments in each fragment size bin that have cut sites of both the enzymes on either end. The predicted fragment size distributions from digestion by different enzyme pairs are plotted. Further outputs are given (by JavaScript code “dataFrame.js”) as an interactive table where the user can try out different size selection cut-offs for each of the enzyme combinations to compare the number of SNPs in the digested sample, the number of samples that can be multiplexed, and the expected sequencing efficiency (see Fig. 1). Sequencing efficiency is calculated as the percentage of sequenced bases that is not wasted on overlaps or adaptor contamination.

In addition to these theoretical values, the table also lists values predicted by models based on empirical data from the meta-analysis and our controlled experiment (see Fig 3 in the Results section), incorporating the effect of incomplete size selection. This is done using two linear models based on a linear regression analysis, one for read overlaps and one for adaptor contamination (Fig. 3; details provided in Methods and Results section). These estimates cannot be taken as precise predictions because of the large variation we observed across the datasets included in the meta-analyses. They are meant to be suggestive of the incomplete effectiveness of size selection and therefore the expected sequencing efficiency. They should only be taken as a cautionary indication of how unrealistic the theoretical predictions might be.

ddgRADer can also be used to visualise in silico digestion by a single enzyme by choosing the same enzyme twice. This might be relevant for methods such as genotyping-by-sequencing (GBS) that involve digestion by a single enzyme. However, it cannot be used for protocols that involve further random shearing of the DNA fragments generated after digestion such as single-digest RAD sequencing. The workflow of ddgRADer is shown in Fig. 1 and full details of the internal calculations and predictions are given in Supp. Table 3. The scripts used in ddgRADer are available on github at https://github.com/felixglinka/ddRadSeqWebTool.

## Materials and methods

### Literature survey

A literature survey was first carried out to review the diversity of study organisms, enzyme pairs and size-selection cut-offs used in ddRADseq studies and estimate the proportion of short fragments resulting from these choices. To keep the number of papers manageable, we limited our search to papers published in Molecular Ecology and Molecular Ecology Resources as these journals routinely publish studies involving ddRADseq. The keywords “ddRAD” and “paired-end” were used for the search in the software Publish or Perish (PoP; Harzing, 2007), which allowed us to get the data in a tabular format. The enzymes used, size-selection method and cut-offs, the stage in which size selection was performed (before or after PCR) and read length were noted. We then downloaded the raw sequencing data from NCBI or dryad and re-analysed them for these studies whenever possible. Most of the published datasets could not be used because of the following reasons: there was no closely related reference genome available for the study species, the sequencing data was not made public, only forward reads were published, filtered sequencing data was published, or mismatched number of reads in the forward and reverse files. We analysed 12 (Baiz et al., 2019; Combs et al., 2018; de Jong et al., 2020; Farleigh et al., 2021; Fritz et al., 2018; Ivanov et al., 2018; Maigret et al., 2020; Portnoy et al., 2015; Ryan et al., 2017; Schley et al., 2020; Termignoni-García et al., 2017; Trense et al., 2021) where size-selection was carried out using Pippin Prep or Bluepippin before the PCR step.

We used FastQC (Andrews, 2010) to estimate the level of adaptor contamination in each of the datasets. Adaptors were then trimmed using Trimmomatic (Bolger et al., 2014) with the default settings, which are to identify a minimum of 8bp adaptor sequence. Sequences were not filtered or trimmed for low quality. FLASH (Magoč & Salzberg, 2011) was used to combine the trimmed overlapping paired-reads and get a fragment size distribution of the combined reads. The output of FLASH was a histogram of all the combined reads that was then used to find the proportion of read overlap and adaptor contamination in the data out of the total number of reads in the empirical datasets. We obtained the theoretically predicted fragment size distribution from the in silico digestion of the closest available reference genome using a custom python scripts (python scripts “HandleFastafile.py” and “DigestSequence.py”, available in the ddgRADer github above). Based on this, we calculated the proportion of fragments (out of total number of fragments shorter than 1000bp) that were below two read lengths for the respective dataset.

To test whether in silico prediction and size-selection criteria significantly predicted the empirical outcome of adaptor contamination and overlapping reads, we performed two multiple linear regressions, one for adaptor contamination (fragments shorter than one read length) and another for read overlaps (between one and two read lengths). The empirical proportion of adaptor contamination (or overlaps) was the response variable, with two predictor variables: in silico proportion of adaptor contamination (or overlaps) and the width of the size-selection range. Note that the in silico proportion is the proportion of short fragments in the in silico digestion of the genome, before any size selection. We also tested for an interaction between these two predictors. Datasets where the size selection cut-off overlapped with the adaptor contamination range or read overlap range were excluded for the respective analyses. This was necessary because the cut-off reported in the papers often did not match with the one observed in the empirical data, and it was difficult to determine the precise cut-off based on visualising the fragment size distributions. We further excluded the study in Supp. Fig. 1l as the empirical results suggested that size-selection failed entirely. This resulted in a total of 12 datasets that were used for the adaptor contamination regression (10 of the 12 datasets from the literature review along with two datasets from our controlled experiment; Supp. Fig. 1l and Fig. 2d were excluded), and eight datasets were used for the read overlap model (eight of the 12 datasets from the literature review; Fig. 2b–f and Supp. Fig. 1l were excluded).

**Figure 2:**
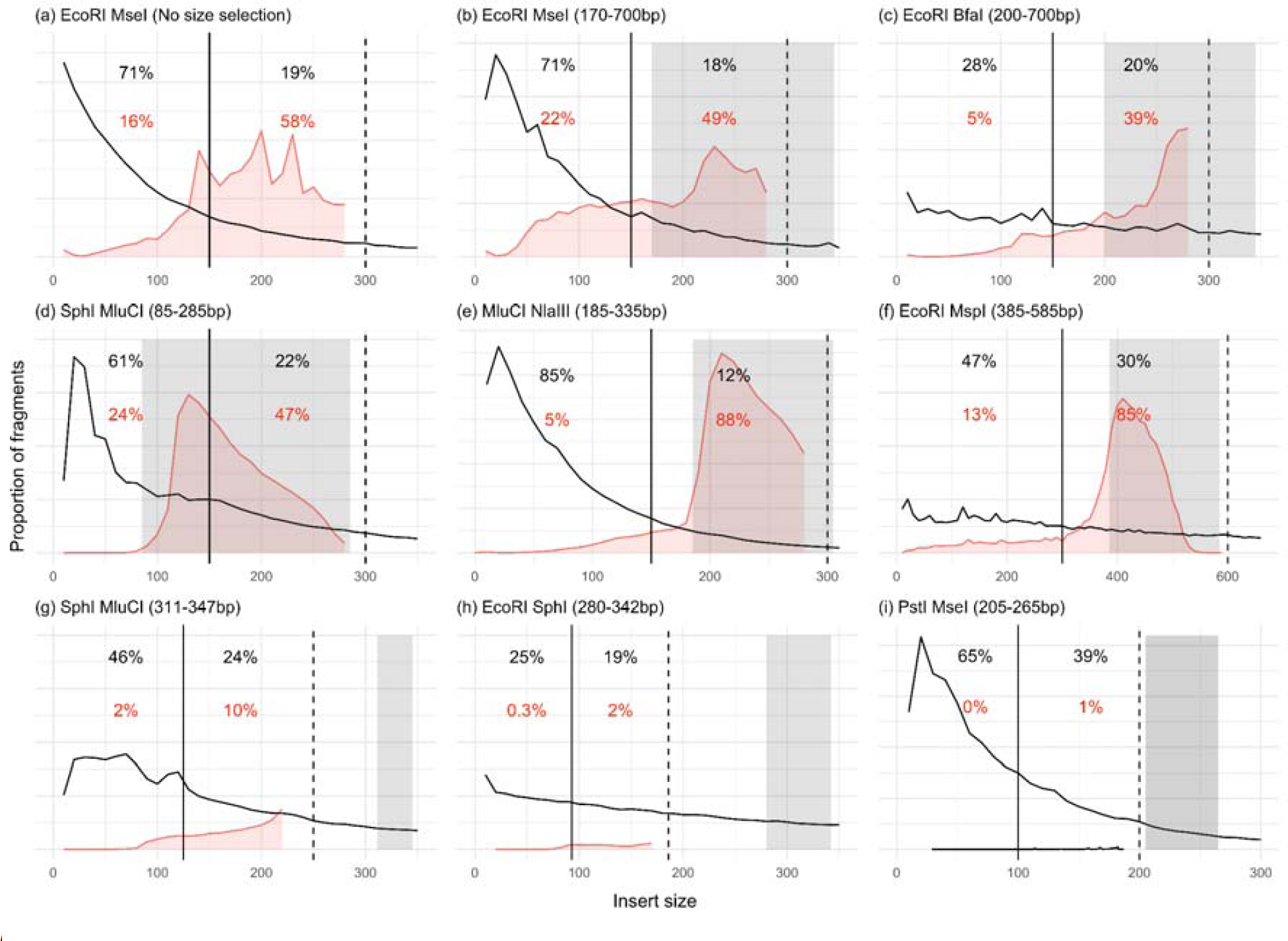
Representative plots of the datasets that were analysed to show in silico and empirical proportion of short fragments. Fragment size distribution from in silico digestion (black trend line) and observed in empirical data (red histogram). Black dashed lines indicate two read lengths, below which overlapping reads are expected. Black solid lines indicate one read length, below which we expect adaptor contamination. Grey box represents the intended insert size selection criteria chosen in the study. Percentages in black represent in silico prediction of the proportion of fragments (before any size selection) and red are the proportion of fragments in empirical data (after size selection). **(a)** Inbar et al., (in prep) did not use Bluepippin for size selection, only magnetic beads clean up, **(b-c)** controlled experiments carried out on the same set of samples while switching the frequently cutting enzyme, **(d-i)** representative studies that used Bluepippin: (d) Baiz et al., 2018 (e) Trense et al., 2020 (f) Fritz et al., 2017 (g) Combs et al., 2017 (h) Termignoni et al., 2017 (i) Ivanov et al., 2018.

### Controlled experiment to study the effect of enzyme choice

We conducted a controlled experiment on *Camponotus* carpenter ants to understand the effect of changing the frequent-cutting enzyme on the proportion of short fragments and consequently on sequencing efficiency.

#### In silico digestion and size-selection cut-offs

The *Camponotus floridanus* reference genome (NCBI accession QANI01000000) was in silico digested to compare the fragment size distributions generated by both pair of enzymes as described above. To estimate the number of genomic sites that were expected to be sequenced at different size-selection cut-offs, taking into account the waste due to adaptor contamination and read overlaps, an excel sheet calculator was created (Supp. File 1). Additionally, we estimated the number of samples that could be multiplexed with a given yield of an Illumina sequencing lane and depth requirement of the study. These calculations were subsequently implemented in the webtool ddgRADer and described in Supp. Table 3.

#### Sampling, library construction, and sequencing

Samples of 125 male *Camponotus fellah* were collected from Rehovot, Israel (31°54’32.1“N 34°48’45.7“E) from a single nest and stored at -80C. DNA was extracted from the abdomen of the ants using the Qiagen DNeasy blood and tissue kit followed by ethanol precipitation of the DNA extracts. Genomic libraries were constructed following Brelsford et al., (2016), a modified double-digest Restriction-site Associated DNA sequencing (ddRADseq) protocol based on Parchman et al. (2012) and Peterson et al. (2012) (Supp. File 2). To understand the effect of enzyme choice on the resulting fragment size distribution and the proportion of waste due to short fragments, we compared libraries built using different restriction enzyme pairs. One way to switch enzymes where the same Illumina sequencing adaptors can be used, is to switch to an enzyme that generates the same 3’ overhang. One set of libraries was constructed using the commonly used combination of EcoRI and MseI. EcoRI is a 6-base rare cutter (recognition sequence 5′-G|AATTC-3′) while MseI is a 4-base frequent cutter (recognition sequence 5′-T|TAA-3′). Given that the recognition site for MseI is GC poor, for a genome with <50% GC content, MseI cuts frequently, generating a large proportion of small fragments. BfaI is a 4-base frequent cutter generating an overhang identical to MseI, yet its recognition site contains more GC than MseI (recognition sequence 5′-C|TAG-3′). Therefore, the GC-poor genome of *C. fellah* contains less BfaI than MseI sites, which should increase the fragment sizes and improve sequencing efficiency (in general, ant genomes contain between 30-50% GC (Simola et al., 2013), and the genome of *C. floridanus* has only 33% GC).

Briefly, genomic DNA was digested using two restriction enzymes and adaptors containing unique barcodes were ligated to each sample for multiplexing (using CutSmart buffer® NEB). The ligated products were PCR amplified in replicates of four using Q5 Hot Start Polymerase® NEB for 20 cycles with a starting DNA volume increased to 7ul per reaction. Additionally, primers and dNTPs were added to a final thermal cycle step to reduce single-stranded or heteroduplex PCR products. All the samples for each pair of enzymes were pooled and the two sets of libraries were size selected using Bluepippin (Sage Science, Beverly, MA, USA). 125 samples were used for MseI/EcoRI and 117 of them were also used for building BfaI/EcoRI libraries. The BfaI/EcoRI libraries were selected for a range of 330 to 700bp and MseI/EcoRI for a range of 300bp to 700 bp (this size includes 128bp of the adaptors and primers). The rationale behind choosing different size-selection ranges are discussed in detail in the Results and discussion section below. The libraries produced with the two sets of enzymes were sequenced with paired end 150bp reads on a single lane of an illumina Hiseq X sequencer to generate 340 million paired end reads (NCBI SRA Accession: xxxxxx).

### ddRADseq data analyses

The quality of the raw sequence data was examined using FastQC. The mean phred quality score was 37.9 and more than 90% of the reads had a mean score >= 30. The total number of paired-reads in the BfaI/EcoRI raw data were 143,893,548 (117 samples) while in MseI/EcoRI were 143,444,471 (125 samples). We removed the excess samples from the MseI/EcoRI dataset to have a comparable set of samples in the two datasets. The BfaI raw data was then subsampled (randomly excluded reads) to have a comparable number of reads in both the datasets. These two datasets were then analysed similar to literature survey datasets using Trimmomatic and FLASH. The command process_radtags.pl from the Stacks2 pipeline (Rochette et al., 2019) was used to demultiplex the data. The resulting reads were mapped to the *Camponotus floridanus* genome (NCBI accession number: GCA_003227725.1 (Shields et al., 2018)) using *Bowtie2* (Langmead B & Salzberg SL, 2012). Using Ref_map.pl from stacks2 the mapped sequences were further processed to call SNPs. The details on the number of reads at each step are given in Supp. Table 1.

## Results and discussion

Adaptor contamination and read overlaps are major factors reducing the efficiency of sequencing. For example, in a previous study in our lab (Inbar et al., in prep) we constructed ddRADseq libraries using samples of the ant *Cataglyphis niger*, with the enzyme combination EcoRI/MseI. This enzyme choice resulted in read overlaps in 74% of read pairs (fragments shorter than two read lengths), of which 16% had adaptor contamination (fragments shorter than one read length; see Fig. 2a). The fact that we only see adaptor contamination in 16% was due to the bead clean-up step in the library construction protocol, which excludes a significant proportion of short fragments. And yet, 27% of the sequencing yield in this study was wasted on adaptor contamination and read overlaps. Another way to put it is that the sequencing efficiency was only 73% in this study, where efficiency is defined as the percentage of sequenced bases that is not wasted on overlaps or adaptor contamination. Many researchers try to deal with this issue using a more precise size selection method, such as manually cutting fragments of a certain size range from a gel or using instruments like Pippin Prep, to exclude short fragments from the library.

We carried out a literature survey to understand the range of size selection criteria and commonly used enzymes in ddRADseq protocols. We reviewed a total of 66 studies published in the journals Molecular Ecology and Molecular Ecology Resources in the years 2015 - 2021 where ddRADseq data were generated using paired-end sequencing (Supp. File 3). Of these, 38 studies (58%) used some form of automated gel electrophoresis-based size selection such as Bluepippin or Pippin Prep, 10 studies (15%) used only magnetic beads for size selection (nine Ampure XP and one Sera-mag magnetic beads), five studies (8%) used manual size selection from a gel, one study used Chroma spin column and 12 studies (20%) did not report the size selection method used. 10 studies (15%) chose a size selection range with the lower cut-off below two read lengths, intentionally including overlapping read pairs. Most studies used a broad size selection range that was wider than 100bp (40 studies: 61%). Among the enzyme pairs, 33 studies (50%) used a combination of 6-base and 4-base cutter, 14 studies (21%) used 8-base and 4-base cutter, 10 studies (15%) used two 6-base cutters, 7 studies (11%) used two 4-base cutters and 2 studies used 8-base and 6-base cutter.

EcoRI/MspI was the most used enzyme pair (10), followed by EcoRI/SphI (8), MluCI/NlaIII (6), SbfI/MseI (6), SbfI/MspI (6), and SphI/MluCI (5) (Supp. Table 2). We then conducted a meta-analysis, reanalysing and comparing 12 ddRADseq datasets from diverse organisms to evaluate the extent of adaptor contamination and read overlaps. These 12 studies generated highly diverse datasets demonstrating a range of possible outcomes from ddRADseq (see representatives in Fig. 2d-i; full results in Supp. Fig. 1), which depend on a combination of factors in the experimental design, including the genome of the study organism, the choice of restriction enzymes, size selection method and cut-offs, and other technical variations in the protocol. Therefore, we conducted a controlled comparative study to examine the effect of alternative enzyme choices on the resulting adaptor contamination, overlapping reads, and consequently, efficiency of sequencing. We generated two sets of libraries for the same set of samples with two different frequently cutting enzymes. Both sets of libraries were built using the same rare cutter (EcoRI), but the frequent cutter (MseI) was switched to an enzyme with a more GC rich recognition site (BfaI), because of which it cut the genome less frequently. Thereby, we increased the average fragment size in the digestion products (Fig. 2b-c).

The first major factor differentiating all these studies was the in silico predicted distribution of fragment sizes. The results of the meta-analyses and our controlled experiment revealed high variation in the shape of the in silico distributions, dramatically differing in their skewness. Some studies had most of the expected distribution in the size range of one read length, which would result in high levels of adaptor contamination. For example, Fig. 2e (using NlaIII/MluCI) and Fig. 2b (our EcoRI/MseI experiment) had 85% and 71% of the in silico distribution below one read length, respectively. In contrast, Fig. 2h (using EcoRI/SphI) and Fig. 2c (our EcoRI/BfaI experiment) had 25% and 28%, respectively. These results show that the initial fragment size distribution, before any size selection, is highly dependent on the enzyme choice.

We further observed considerable variation in the efficacy of size selection in eliminating short fragments. In theory, one might expect no adaptor contamination and read overlaps if the size selection cut-off is larger than two read lengths. However, in practice we observed large proportion of sequencing reads with overlap and adaptor contamination in some of the studies. For example, the study in Fig. 2g chose a size selection cut-off that was bigger than two read lengths, and yet 10% of the sequencing output had read overlaps and 2.1% had adaptor contamination. Some studies intentionally allowed read overlaps but had a size selection cut-off just bigger than one read length. In these cases, although adaptor contamination was not expected, large proportions of adaptors were observed (e.g., 13% in Fig. 2f). These results demonstrate that a large proportion of short fragments might pass through the size selection procedure, even when using the most precise instruments available. This issue is worse when the in silico distribution is more skewed, i.e., having a larger proportion of short fragments before size selection. This effect explains the difference observed in our controlled experiment, with 22% adaptor contamination in Fig. 2b (EcoRI/MseI) and only 5% in Fig. 2c (EcoRI/BfaI), corresponding to the difference in their in silico proportions of 71% and 28%, respectively.

The second major factor differentiating ddRADseq studies was the width of the size selection window (ranging from 36 to 520bp). Half of the studies used a narrow size selection range (the minimum width possible under “tight mode”, which corresponds to a target size ±16% according to the 2018 Bluepippin manual). Unlike the studies that used a broad range, this approach effectively resulted in negligible waste of sequencing due to read overlap and adaptor contamination (Fig. 2g, 2h, 2i, Supp. Fig. 1k, 1n, 1o). However, its drawback was that a relatively small number of genomic loci were sequenced, typically resulting in a large proportion of sequence duplication, which is another kind of waste (data not shown).

Size selection not only leads to variable outcomes, but it also appears to be imprecise. In our controlled experiments (Fig. 2b and 2c), we see a shift of about 50bp between the intended size selection range and the outcome in the empirical data. A similar shift of about 40bp can be observed in Fig. 2d. Despite these shortcomings of size selection, we see a considerable effect of size selection in enriching fragments within the desired range. For studies shown in Fig. 2b–f, size selection cut-offs were below two read lengths, which allows us to observe the proportion of the empirical distribution within the size selection range. Size selection in these studies resulted in a much larger proportion of fragments in the selected range, higher than that expected based on the in silico distribution, indicating that size selection was successful in enriching target fragment sizes.

We also observed an effect of the distance between the size selection cut-off and the read length on the efficacy of size selection. For example, both studies in Fig. 2f and Fig. 2g had a similar proportion of predicted fragments below one read length (46% vs 47%). In Fig. 2g, the size selection range begins above two read lengths while in Fig. 2f the lower cut-off is just larger than one read length. This resulted in only 2% adaptor contamination in Fig. 2g compared to 13% in Fig. 2f, suggesting that as the lower cut-off comes closer to the one read length line, adaptor contamination increases. This might be due to the tendency of some short fragments to migrate with slightly longer fragments in gel electrophoresis. However, choosing a high selection cut-off might not necessarily be the optimal choice, because it might dramatically reduce the number of loci in the sequencing output.

We formulated a mathematical model based on these insights. We carried out multiple linear regression analyses to predict empirical outcome of adaptor contamination (Fig. 3a) and read overlaps (Fig. 3b) as a function of the in silico digestion (predictor 1) and the width of the size-selection range (predictor 2). These analyses regress the proportions observed in the sequencing results after size selection against the proportions in the in silico digestion without any size selection. Thereby, we can see if highly skewed in silico distributions with a large proportion of short fragments result in a large proportion of short fragments that fail to be excluded by size selection. For adaptor contamination, the fitted model was:

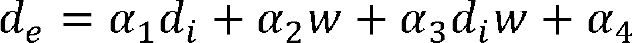

**Figure 3:**
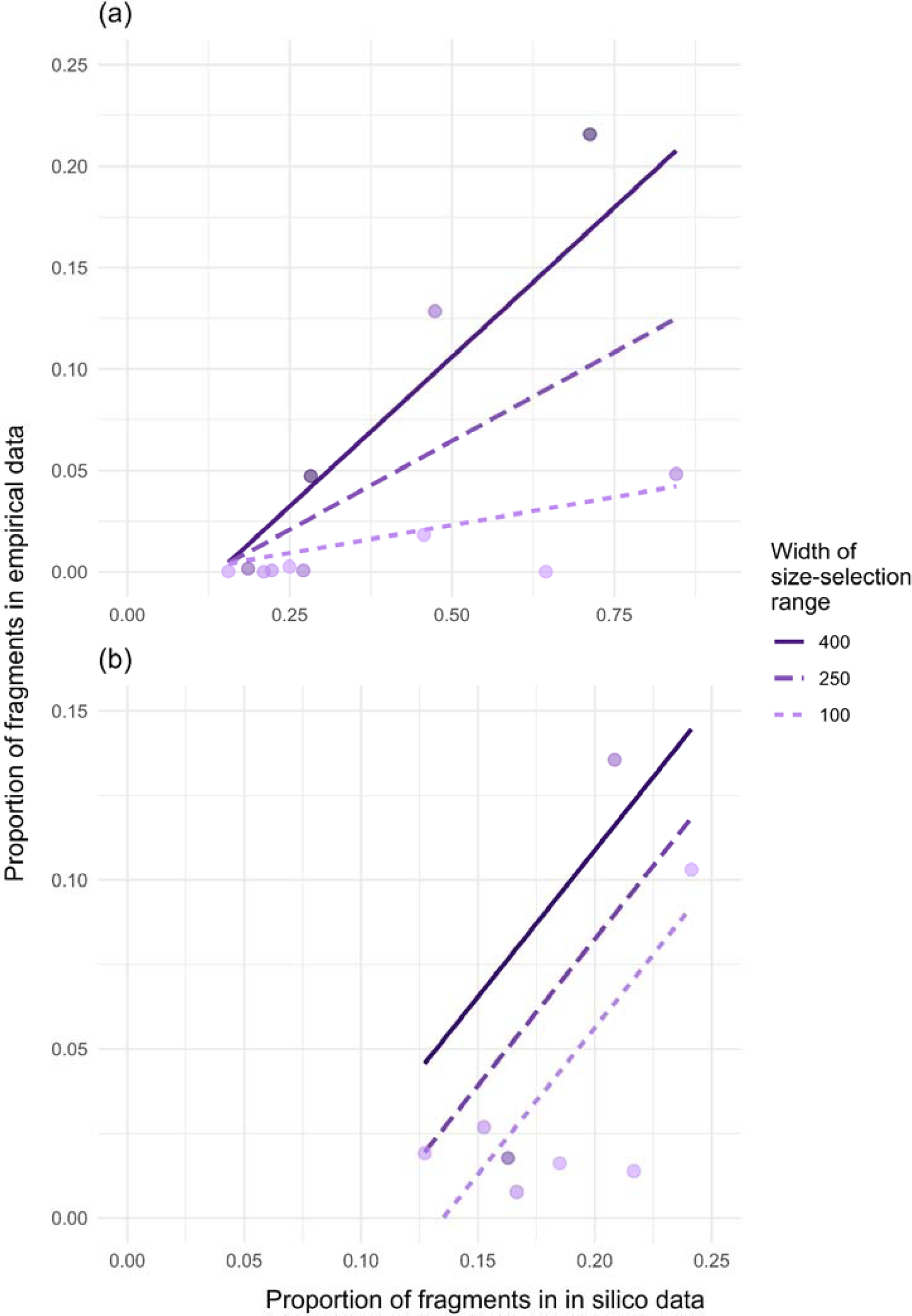
Linear models to predict the proportions of (a) adaptor contamination (fragments shorter than one read length) and (b) read overlaps (fragments longer than one and shorter than two read lengths). Empirical proportions of adaptor contamination and read overlaps observed in the analysed datasets were plotted as a function of in silico proportions of adaptor contamination and read overlaps. Regression lines are plotted for three representative widths of the size-selection range.

where *d_e_* is the empirical proportion of adaptor contamination, *d_i_* is the in silico proportion of adaptor contamination, *w* is the width of the size selection range, and the coefficients are: L*_1_*= –0.0245421, L*_2_*= –0.0001226, L*_3_*= 0.0007978, L*_4_*=0.0077016. The regression was statistically significant (F-statistic: 13.96, DF: 8, p-value: 0.0015, Adjusted R-squared: 0.7795). Note that the interaction term (L*_3_d_i_w*) results in a substantial change in the slope of the regression line as the size-selection range becomes wider (see three representative regression lines in Fig. 3a). This model suggests that the in silico distribution and width of the size-selection range together have a large effect on empirical outcome of adaptor contamination. However, the multiple regression model for read overlaps (Fig. 3b) was not significant with the interaction term (F-statistic: 1.956; DF: 4, p-value: 0.26, Adjusted R-squared: 0.2907). Consequently, we dropped the interaction term from the model to get an additive model. This model was also not significant (F-statistic: 1.878 on 2 and 5 DF, p-value: 0.25, Adjusted R-squared: 0.2005). The multiple regression additive model for overlapping reads was:

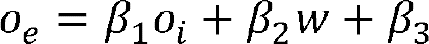

where *o_e_* is the empirical proportion of overlaps, *o_i_* is the in silico proportion of overlaps, *w* is the width of the size selection range, and the coefficients are: β*_1_*= 0.8672884, β*_2_*= 0.0001751, β*_3_*= –0.1347334. These two models for predicting the proportions of adaptor contamination and read overlaps were incorporated into ddgRADer to give the user a more realistic prediction of the problem of short fragments after size selection. The detailed summary table of the models are provided in Supp. Table 4 and 5. Although the model for predicting overlapping reads was not significant, we implemented it in ddgRADer to give the user some idea about the scale of overlaps that can be expected. The webtool also displays a disclaimer to exercise caution while relying on these estimates.

One of the differences between our two controlled experiments, other than the frequently cutting enzyme, was the size selection cut-off. In the EcoRI/MseI experiment, the in-silico distribution predicted close to 90% of the fragments to be below 300bp compared to only 48% in EcoRI/BfaI. While selecting a cut-off above 300bp would result in efficient sequencing with low wastage, this would also exclude ∼70% of the genomic sites from being sequenced (only 5 million out of a potential 18 million genomic sites would be sequenced for EcoRI/MseI; Fig. 4). Therefore, we considered the trade-off between genomic sites sequenced and sequencing efficiency, defined as the proportion of sequenced bases not wasted due to adaptor contamination or read overlaps (see Methods for details). As the lower cut-off moves below 300bp, we allow some read overlaps and sequencing efficiency drops. This drop is more pronounced for MseI than BfaI (dashed lines in Fig. 4) in line with their different in silico distributions. Hence, different cut-offs were chosen for the two experiments to optimise this trade-off: 170bp for EcoRI/MseI and 200bp for EcoRI/BfaI.

**Figure 4:**
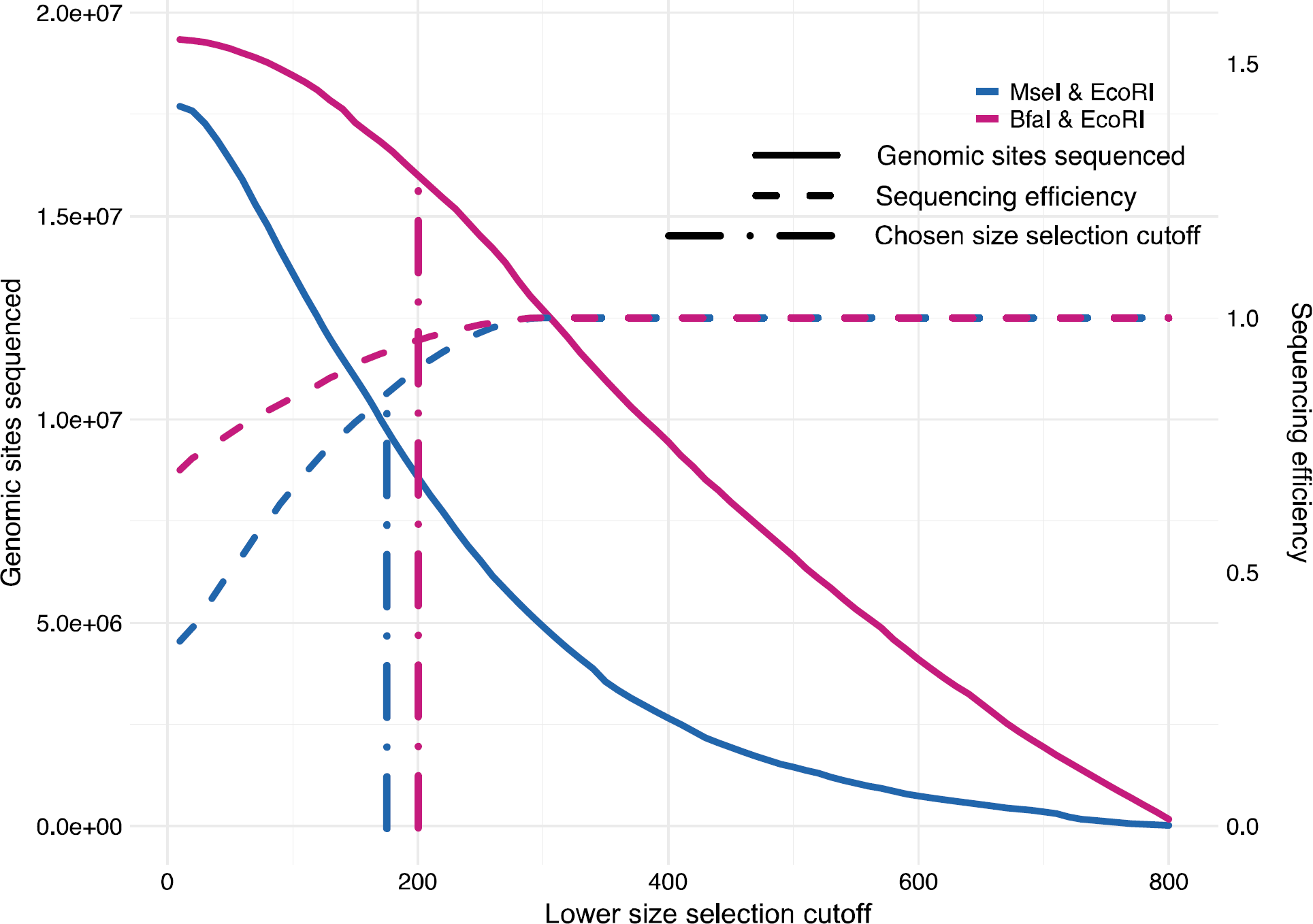
Trade-off between sequencing efficiency and the number of genomic sites sequenced plotted against the size selection cut-off for the controlled experiment.

Theoretically, we expected an efficiency of 92% in EcoRI/MseI (for the observed sharp drop in the empirical distribution at 210bp in Fig. 2g) and 98% in EcoRI/BfaI (at 240bp in Fig. 2h) based on the excel sheet calculator (Supp. File 1). However, due to poor efficiency of size selection that allows unintended shorter fragments to go through, we observed lower sequencing efficiency. Moreover, the considerably lower sequencing efficiency of EcoRI/MseI relative to EcoRI/BfaI is a reflection of the larger number of short fragments produced by EcoRI/MseI compared to EcoRI/BfaI as predicted from the in silico distribution. The empirical data from the controlled experiment revealed a sequencing waste of 27% (sequencing efficiency of 73%) in EcoRI/MseI compared to 2.7% in EcoRI/BfaI (sequencing efficiency of 97%) as a result of adaptor contamination and read overlaps.

Interestingly, we observe higher variance of sequencing depth for EcoRI/MseI compared to EcoRI/BfaI (Fig 5a). Accordingly, if we do not set any threshold on the minimum depth, we have 8,003,694 unique genomic sites sequenced with EcoRI/MseI compared to 5,404,131 with EcoRI/BfaI. However, as in common practice, where we do require a minimum depth of 10X (the --minDP parameter in vcftools), less genomic sites are usable in EcoRI/MseI (513,891) compared to EcoRI/BfaI (737,766) (Fig. 5b). Therefore, the enzyme choices that give a more uniform fragment size distribution ultimately result in a larger number of genomic sites that can be genotyped (for a given sequencing budget, and with a commonly used depth cut-off).

**Figure 5:**
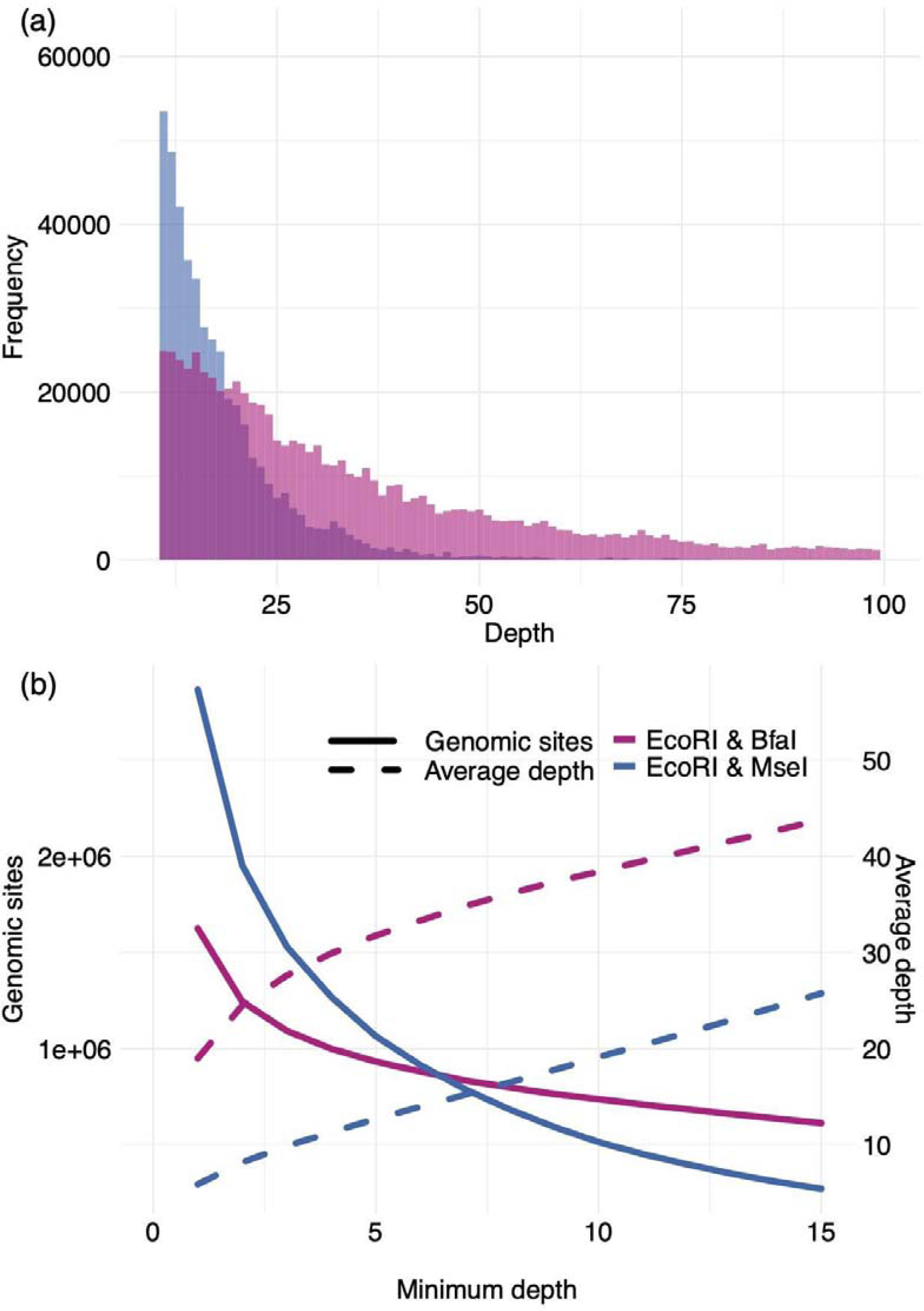
(a) Distribution of sequencing depth for the two enzyme choices. **(b)** Number of unique genomic sites sequenced plotted against the minimum depth cut-off.

The insights from the above analyses were implemented in the webtool ddgRADer to aid researchers in optimising ddRADseq experimental design (described in the beginning of this manuscript). A few previously published tools were already available for researchers. SimRAD (Lepais & Weir, 2014) is designed to choose between different reduced representation library protocols by predicting the number of loci that can be generated. It performs in silico digestion and fragment size-selection allowing the user to choose the enzyme and size-selection criteria. Another software that carries out a similar analysis is DDRADSEQTOOLS (Mora-Márquez et al., 2017). In addition to in silico digestion, this tool also simulates the effect of allele dropout and PCR duplicates on the coverage taking into account potential sources of error. Both simRAD and DDRADSEQTOOLS are R packages. RADinitio (Rivera-Colón et al., 2021) is a python software that simulates population-level data based on a demographic model input by the user over a reference sequence. The user can implement various reduced representation protocols, in silico digestion, size selection, library amplification, and sequencing depth, while considering errors such as allelic dropout. Additionally, RADinitio carries out retrospective simulation, where an in silico library is generated based on an empirical RADseq dataset. Thus, RADinitio takes into account a lot more variables, particularly inherent variability in natural populations, that contribute to the quality of the data. While some of these tools help optimise experiments by taking into account allele dropouts and PCR duplication, none of them help in making the experiment cost effective by optimising the sequencing efficiency. Moreover, they assume perfect size-selection and do not take into account its incomplete effectiveness, which results in additional adaptor contamination and read overlaps. This is an important factor to consider when selecting the enzyme pair because in certain cases it has a dramatic effect on sequencing efficiency, the expected number of SNPs, the sequencing depth, and the number of samples that can be multiplexed per lane. Additionally, all these softwares are command-line tools and might require some skill in R, which would involve a steep learning curve particularly for researchers without prior experience with command-based programs.

The easy-to-use graphical user interface of ddgRADer makes it accessible to even a beginner, considerably reducing the time taken to design ddRADseq experiments. Further, multiple popovers guide the user through the process and additional documentation is included to help in decision making. Thereby, this tool can aid researchers that do not yet have experience or in-depth understanding of ddRADseq protocols and the associated issues. ddgRADer explicitly incorporates the error associated with size-selection by accounting for additional shorter fragments that result in adaptor contamination and read overlaps. This helps the user make informed enzyme choices taking into account sequencing efficiency. Although enzymes aren’t very expensive, synthesising the multiplexing barcodes requires a large initial investment. These barcodes attach to the overhang generated by the restriction enzyme making enzyme choice a consequential decision. Additionally, ddgRADer allows the user to compare several enzyme pairs and the size-selection cut-offs that best suit each of these pairs. The interactive size-selection slide bar helps the user understand the trade-off between the number of genomic sites in the digested sample and sequencing efficiency, as a function of different size-selection cut-offs.

## Conclusion

Here, we demonstrate using empirical data and controlled experiments that enzyme choice greatly affects inadvertent inclusion of adaptor contamination and read overlaps, in spite of size-selection. We observed that the efficacy of size-selection is variable across studies and imprecise. Therefore, enzyme choice is crucial in determining sequencing efficiency and optimising ddRADseq experimental design. We provide a user-friendly webtool that guides beginners through considering alternative enzyme choices and optimising size-selection. The tool provides realistic predictions about the expected outcomes from ddRADseq, based on our empirical characterization of size-selection efficacy in a meta-analysis across diverse studies from the literature as well as our controlled experiments. This will make ddRADseq far more accessible and facilitate the smooth transition of more researchers into the field of population genomics.

## Acknowledgements

We thank Jessica Purcell and Alan Brelsford for advice on ddRADseq methodology, Pnina Cohen for help in sample collection, and Viraj Torsekar for discussions and advice on statistical analyses. AL thanks Praveen Karanth for providing lab space. We are grateful to the reviewers for their insightful comments that improved the manuscript considerably. This study was funded by US-Israel Binational Science Foundation (Grant no. 2017319).

## Data Accessibility Statement

Genetic data:

Raw sequence reads will be deposited in the Sequence Read Archive (NCBI SRA) upon acceptance (BioProject XXX)

## Benefit-Sharing Statement

This is not applicable.

## Notes

### Competing Interest Statement

The authors have declared no competing interest.

### Summary of Updates

We have added additional datasets from literature survey that have now considerably improved the predictions of our model. These models are now explained in greater detail. We have also added a link to the github page.

## References

Andrews, S. (2010). FastQC: A quality control tool for high throughput sequence data. Babraham Bioinformatics, *Babraham Institute*, Cambridge, United Kingdom.

Andrews, K. R., Good, J. M., Miller, M. R., Luikart, G., & Hohenlohe, P. A. (2016). Harnessing the power of RADseq for ecological and evolutionary genomics. Nature Reviews Genetics, 17(2), 81–92. https://doi.org/10.1038/nrg.2015.28

Baird, N. A., Etter, P. D., Atwood, T. S., Currey, M. C., Shiver, A. L., Lewis, Z. A., Selker, E. U., Cresko, W. A., & Johnson, E. A. (2008). Rapid SNP Discovery and Genetic Mapping Using Sequenced RAD Markers. PLoS ONE, 3(10), e3376.

Baiz, M. D., Tucker, P. K., & Cortés-Ortiz, L. (2019). Multiple forms of selection shape reproductive isolation in a primate hybrid zone. Molecular Ecology, 28(5), 1056–1069. https://doi.org/10.1111/mec.14966

Bolger, A. M., Lohse, M., & Usadel, B. (2014). Trimmomatic: A flexible trimmer for Illumina sequence data. Bioinformatics, 30(15), 2114–2120.

Brelsford, A., Dufresnes, C., & Perrin, N. (2016). High-density sex-specific linkage maps of a European tree frog (Hyla arborea) identify the sex chromosome without information on offspring sex. Heredity, 116(2), Article 2. https://doi.org/10.1038/hdy.2015.83

Clugston, J. A. R., Kenicer, G. J., Milne, R., Overcast, I., Wilson, T. C., & Nagalingum, N. S. (2019). RADseq as a valuable tool for plants with large genomes—A case study in cycads. Molecular Ecology Resources, 19(6), 1610–1622.

Catchen, J., Hohenlohe, P. A., Bassham, S., Amores, A., & Cresko, W. A. (2013). Stacks: An analysis tool set for population genomics. Molecular Ecology, 22(11), 3124–3140.

Catchen, J. M., Amores, A., Hohenlohe, P., Cresko, W., & Postlethwait, J. H. (2011). *Stacks*: Building and Genotyping Loci *De Novo* From Short-Read Sequences. G3 Genes|Genomes|Genetics, 1(3), 171–182.

Combs, M., Puckett, E. E., Richardson, J., Mims, D., & Munshi-South, J. (2018). Spatial population genomics of the brown rat (Rattus norvegicus) in New York City. Molecular Ecology, 27(1), 83– 98.

Davey, J. W., Hohenlohe, P. A., Etter, P. D., Boone, J. Q., Catchen, J. M., & Blaxter, M. L. (2011). Genome-wide genetic marker discovery and genotyping using next-generation sequencing. Nature Reviews Genetics, 12(7), 499–510.

de Jong, M. J., Li, Z., Qin, Y., Quéméré, E., Baker, K., Wang, W., & Hoelzel, A. R. (2020). Demography and adaptation promoting evolutionary transitions in a mammalian genus that diversified during the Pleistocene. Molecular Ecology, 29(15), 2777–2792.

Eaton, D. A. R., & Ree, R. H. (2013). Inferring Phylogeny and Introgression using RADseq Data: An Example from Flowering Plants (Pedicularis: Orobanchaceae). Systematic Biology, 62(5), 689–706.

Eaton, D. A. R., & Overcast, I. (2020). ipyrad: Interactive assembly and analysis of RADseq datasets. Bioinformatics, 36(8), 2592–2594.

Elshire, R. J., Glaubitz, J. C., Sun, Q., Poland, J. A., Kawamoto, K., Buckler, E. S., & Mitchell, S. E. (2011). A Robust, Simple Genotyping-by-Sequencing (GBS) Approach for High Diversity Species. PLOS ONE, 6(5), e19379.

Farleigh, K., Vladimirova, S. A., Blair, C., Bracken, J. T., Koochekian, N., Schield, D. R., Card, D. C., Finger, N., Henault, J., Leaché, A. D., Castoe, T. A., & Jezkova, T. (2021). The effects of climate and demographic history in shaping genomic variation across populations of the Desert Horned Lizard (Phrynosoma platyrhinos). Molecular Ecology, 30(18), 4481–4496.

Fritz, M. L., DeYonke, A. M., Papanicolaou, A., Micinski, S., Westbrook, J., & Gould, F. (2018). Contemporary evolution of a Lepidopteran species, Heliothis virescens, in response to modern agricultural practices. Molecular Ecology, 27(1), 167–181.

Harzing, A. W. (2007). Publish or perish 6.

Inbar, S., Cohen, P., Yahav, T., & Privman, E. (2020). Comparative study of population genomic approaches for mapping colony-level traits. PLoS Computational Biology, 16(3), e1007653.a

Inbar, S., Saied, B., Cohen, P., Frenkel, Z., Yahav, T., Korol, A., & Privman, E. (In prep). Genetic architecture of nestmate recognition cues.

Ivanov, V., Lee, K. M., & Mutanen, M. (2018). Mitonuclear discordance in wolf spiders: Genomic evidence for species integrity and introgression. Molecular Ecology, 27(7), 1681–1695. https://doi.org/10.1111/mec.14564

Lajmi, Glinka and Privman 2022; ddRADseq data for Camponotus fellah; National Centre for Biotechnology Information: Sequence Read Archive (NCBI SRA); [dataset]

Langmead B, & Salzberg SL. (2012). Fast gapped-read alignment with Bowtie 2. Nature Methods, 9, 357– 359.

Lepais, O., & Weir, J. T. (2014). SimRAD: An R package for simulation-based prediction of the number of loci expected in RADseq and similar genotyping by sequencing approaches. Molecular Ecology Resources, 14(6), 1314–1321.

Luikart, G., England, P. R., Tallmon, D., Jordan, S., & Taberlet, P. (2003). The power and promise of population genomics: From genotyping to genome typing. Nature Reviews Genetics, 4(12), 981– 994.

Magalhaes, I. S., Whiting, J. R., D’Agostino, D., Hohenlohe, P. A., Mahmud, M., Bell, M. A., Skúlason, S., & MacColl, A. D. C. (2021). Intercontinental genomic parallelism in multiple three-spined stickleback adaptive radiations. Nature Ecology & Evolution, 5(2), 251–261.

Magoč, T., & Salzberg, S. L. (2011). FLASH: Fast length adjustment of short reads to improve genome assemblies. Bioinformatics, 27(21), 2957–2963.

Maigret, T. A., Cox, J. J., & Weisrock, D. W. (2020). A spatial genomic approach identifies time lags and historical barriers to gene flow in a rapidly fragmenting Appalachian landscape. Molecular Ecology, 29(4), 673–685.

Mora-Márquez, F., García-Olivares, V., Emerson, B. C., & López de Heredia, U. (2017). ddRADseqTools: A software package for in silico simulation and testing of double-digest RADseq experiments. Molecular Ecology Resources, 17(2), 230–246.

Nadukkalam Ravindran, P., Bentzen, P., Bradbury, I. R., & Beiko, R. G. (2019). RADProc: A computationally efficient de novo locus assembler for population studies using RADseq data. Molecular Ecology Resources, 19(1), 272–282.

Natesh, M., Atla, G., Nigam, P., Jhala, Y. V., Zachariah, A., Borthakur, U., & Ramakrishnan, U. (2017). Conservation priorities for endangered Indian tigers through a genomic lens. Scientific Reports, 7(1), 9614.

Parchman, T. L., Gompert, Z., Mudge, J., Schilkey, F. D., Benkman, C. W., & Buerkle, C. A. (2012). Genome-wide association genetics of an adaptive trait in lodgepole pine. Molecular Ecology, 21(12), 2991–3005.

Peterson, B. K., Weber, J. N., Kay, E. H., Fisher, H. S., & Hoekstra, H. E. (2012). Double Digest RADseq: An Inexpensive Method for De Novo SNP Discovery and Genotyping in Model and Non-Model Species. PLoS ONE, 7(5), e37135.

Portnoy, D. S., Puritz, J. B., Hollenbeck, C. M., Gelsleichter, J., Chapman, D., & Gold, J. R. (2015). Selection and sex-biased dispersal in a coastal shark: The influence of philopatry on adaptive variation. Molecular Ecology, 24(23), 5877–5885.

Puritz, J. B., Hollenbeck, C. M., & Gold, J. R. (2014). *dDocent*: A RADseq, variant-calling pipeline designed for population genomics of non-model organisms. PeerJ, 2, e431.

Rivera[Colón, A. G., Rochette, N. C., & Catchen, J. M. (2021). Simulation with RADinitio improves RADseq experimental design and sheds light on sources of missing data. Molecular Ecology Resources, 21(2), 363–378.

Rochette, N. C., Rivera[Colón, A. G., & Catchen, J. M. (2019). Stacks 2: Analytical methods for paired[end sequencing improve RADseq[based population genomics. Molecular Ecology, 28(21), 4737–4754.

Ryan, S. F., Fontaine, M. C., Scriber, J. M., Pfrender, M. E., O’Neil, S. T., & Hellmann, J. J. (2017). Patterns of divergence across the geographic and genomic landscape of a butterfly hybrid zone associated with a climatic gradient. Molecular Ecology, 26(18), 4725–4742.

Schley, R. J., Pennington, R. T., Pérez-Escobar, O. A., Helmstetter, A. J., de la Estrella, M., Larridon, I., Sabino Kikuchi, I. A. B., Barraclough, T. G., Forest, F., & Klitgård, B. (2020). Introgression across evolutionary scales suggests reticulation contributes to Amazonian tree diversity. Molecular Ecology, 29(21), 4170–4185.

Shields, E. J., Sheng, L., Weiner, A. K., Garcia, B. A., & Bonasio, R. (2018). High-Quality Genome Assemblies Reveal Long Non-coding RNAs Expressed in Ant Brains. Cell Reports, 23(10), 3078– 3090.

Simola, D. F., Wissler, L., Donahue, G., Waterhouse, R. M., Helmkampf, M., Roux, J., … & Gadau, J. (2013). Social insect genomes exhibit dramatic evolution in gene composition and regulation while preserving regulatory features linked to sociality. Genome research, 23(8), 1235–1247.

Sovic, M. G., Fries, A. C., & Gibbs, H. L. (2015). AftrRAD: A pipeline for accurate and efficient *de novo* assembly of RADseq data. Molecular Ecology Resources, 15(5), 1163–1171.

Termignoni-García, F., Jaramillo-Correa, J. P., Chablé-Santos, J., Liu, M., Shultz, A. J., Edwards, S. V., & Escalante-Pliego, P. (2017). Genomic footprints of adaptation in a cooperatively breeding tropical bird across a vegetation gradient. Molecular Ecology, 26(17), 4483–4496.

Toonen, R. J., Puritz, J. B., Forsman, Z. H., Whitney, J. L., Fernandez-Silva, I., Andrews, K. R., & Bird, C. E. (2013). ezRAD: A simplified method for genomic genotyping in non-model organisms. PeerJ, 1, e203. https://doi.org/10.7717/peerj.203

Trense, D., Schmidt, T. L., Yang, Q., Chung, J., Hoffmann, A. A., & Fischer, K. (2021). Anthropogenic and natural barriers affect genetic connectivity in an Alpine butterfly. Molecular Ecology, 30(1), 114–130.

Wagner, C. E., Keller, I., Wittwer, S., Selz, O. M., Mwaiko, S., Greuter, L., Sivasundar, A., & Seehausen, O. (2013). Genome-wide RAD sequence data provide unprecedented resolution of species boundaries and relationships in the Lake Victoria cichlid adaptive radiation. Molecular Ecology, 22(3), 787–798.

Zecherle, L. J., Nichols, H. J., Bar-David, S., Brown, R. P., Hipperson, H., Horsburgh, G. J., & Templeton, A. R. (2021). Subspecies hybridization as a potential conservation tool in species reintroductions. Evolutionary Applications, 14(5), 1216–1224.

