## Supplementary material for "Optimising ddRAD sequencing for population genomic studies with ddgRADer": Supp. File 2

### ddgRADer

The whole application was written using the Django (Version 3.2.4) framework. The user interface was written using JavaScript, CSS and HTML5. A Bootstrap framework (http://getbootstrap.com) was deployed and for the popover windows we used the Popperjs (https://popper.js.org). Backend requests are to be handled using Python (3.9.5). Internally calculations were performed with the help of the pandas package (McKinney, W. et al., 2010) and numpy package (Harris, C.R. et al., 2020). Graphs displayed on the user interface were produced using the Matplotlib (Hunter, J.D., 2007). Furthermore the uploaded fasta file was processed with the help of the Biopython package (Cock, P.J. et al., 2009). The provided restriction enzymes were taken by the New England BioLabs Inc (https://international.neb.com/tools-and-resources/selection-charts/isoschizomers, accessed on 03.06.2021). The web server tool was tested using Microsoft Windows 10 Pro Version 10.0.19042 using Chrome Version 92.0.4515.131 and Firefox Version 90.0.2.

### Workflow outline

The user begins by uploading a relevant genome of their choice. Based on their expertise or requirement, the user can choose enzyme pairs from a list (advanced) or let ddgRADer recommend enzyme options from a predetermined set of commonly used enzyme pairs (beginner). The commonly used rare cutter in the literature were EcoRI, PstI, SbfI and SphI, and for each of these the following frequent cutters are used— EcoRI: MseI, MspI, SphI, SbfI and NlaIII; for PstI: MspI, SphI, MseI, HhaI; for SbfI: EcoRI, MseI, MspI, SphI; and for SphI: MluCI. In the beginner option, depending on the nature of the study design, the user has to specify the expected SNP density along with either the required number of SNPs (for example in a demographic study) or the SNP density to be genotyped (in a study involving genomic scan).

Other parameters that are entered by the user which are common to both the beginner and advanced settings include sequencing read length, sequencing yield, expected SNP density and the desired depth. For each enzyme pair, the tool counts the number of fragments having cut sites of both the enzymes on either side and bins them into 10bp bins up to a maximum fragment size of 1000bp. The webtool outputs a plot of fragment size distribution by the various enzyme pairs (fig. 5b). This data can also be downloaded as a CSV file. Based on the user defined parameters, theoretical and predicted outcomes are tabulated (fig. 5c). The output includes the following:

1) No. fragments: The total number of fragments that are generated by each enzyme pair
2) No. base pairs in insilico digested sample: the total number of genomic nucleotide positions that will be sequenced. For each 10bp bin, the number of base pairs that will be sequenced is calculated by multiplying the number of fragments with the fragment size (e.g. if the bin of 90–100 contains 10 fragments, the number of base pairs would be 1000). For single end sequencing, the sequenced fragment will be only as long as the read length. Therefore, once the bin size exceeds the read length, the number of fragments will be multiplied by the read length. Similarly, in paired end sequencing the number of fragments will be multiplied by double the read length.
3) No. of SNPs in digestion: the expected number of SNPs that will be genotyped is calculated by multiplying the previously calculated <No. base pairs in insilico digested sample> by the expected SNP density in the genome (entered by the user).
4) No. of samples multiplexable: the number of samples that can be multiplexed is calculated by dividing the sequencing yield (amount of data generated by a single lane of sequencer) by the total number of genomic fragments in the size selected range times the depth:

$$No. of\ samples\ multiplexable = \frac{sequencing\ yield}{total\ number\ of\ genomic\ fragments\ in\ the\ size\ selected\ range\ \times\ desired\ depth}$$

5) Sequencing efficiency: calculated as the ratio of the effective total output (number of basepairs sequenced in all multiplexed samples), divided by the maximal potential output, which is calculated based on the sequencing yield and the desired depth:

$$Sequencing\ efficiency = \frac{No.basepairs\ in\ insilico\ digested\ sample\ x\ No.samples\ multiplexable}{sequencing\ yield\ x\ read\ length\ x\ 2*\ /\ desired\ depth}$$

** for paired end sequencing*

6) Fragments shorter than one read length: Fragments shorter than the read length are expected to carry adaptor contamination.
7) Fragments between one read length and two read lengths: In case of paired end sequencing, the overlaps are calculated, as the sum of all fragments between the size of one read length and twice the read length.

A range slider bar is provided for size-selection that can be used on each enzyme pair independently. For each of these parameters, the theoretical and our predicted values are calculated. Although the theoretical values take adaptor contamination and overlapping reads into account, they do not take into consideration the incomplete efficacy of size-selection. The predicted values are calculated using the two linear models from the multiple linear regression analyses (see Fig. 3 and accompanying text in the Results section). In order to recommend the enzyme options, the sum of the nucleotides of the fragments between a size of 300 to 700 is multiplied by the expected polymorphism (expected number of SNPs per kbp) to get the expected number of SNPs. If the resulting value is smaller than the desired number of SNPs, the enzyme pair is excluded in case of enzymes recommended by the webtool. A maximum of five pairs are presented that are predicted to give close to the desired number of SNPs. In case of enzyme pairs chosen by the user, all the above calculations are made for each enzyme pair.

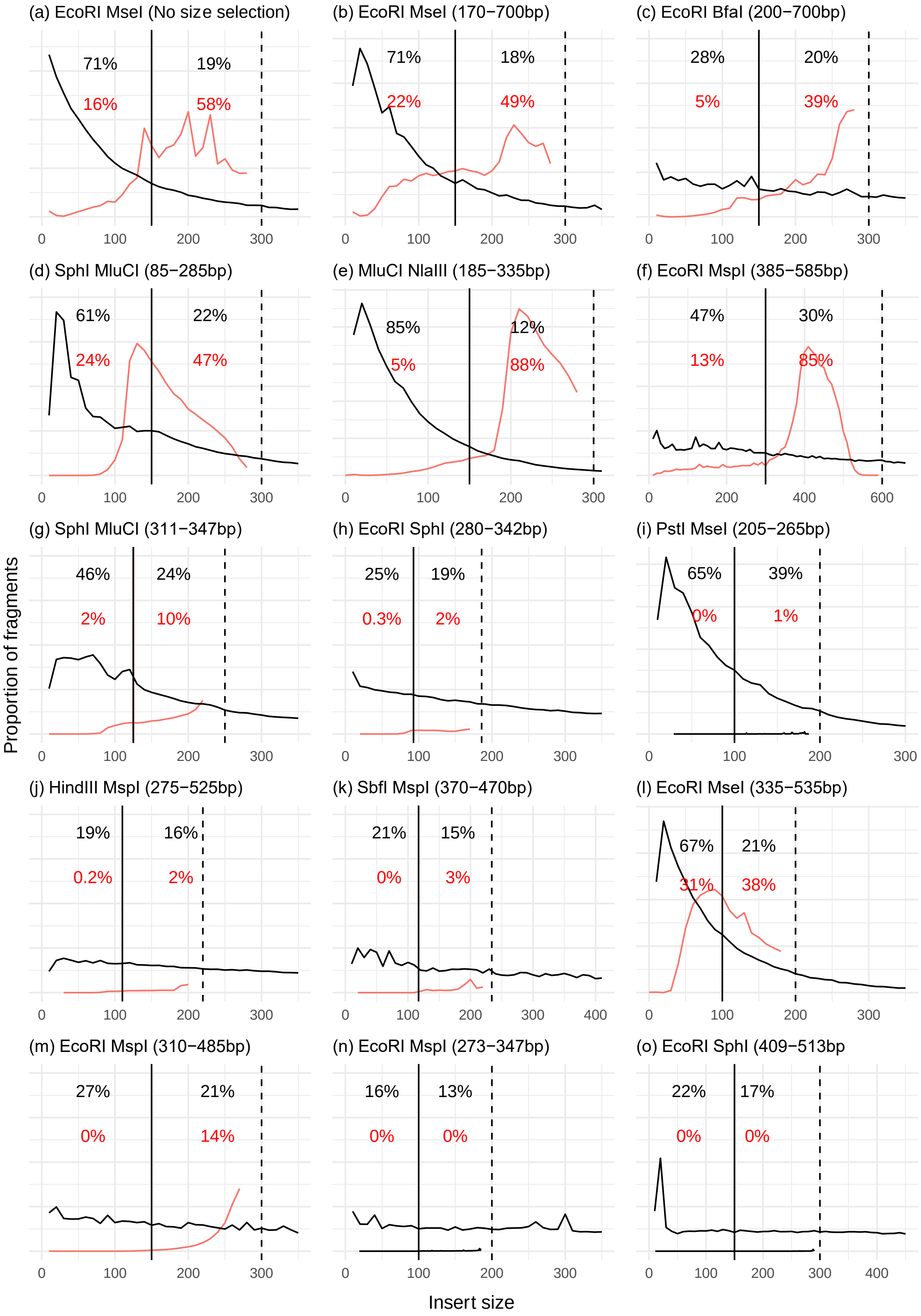

(a) EcoRI MseI (No size selection)
71%
19%
16%
58%
(b) EcoRI MseI (170−700bp)
71%
18%
22%
49%
(c) EcoRI BfaI (200−700bp)
28%
20%
5%
39%
(d) SphI MluCI (85−285bp)
61%
22%
24%
47%
(e) MluCI NlaIII (185−335bp)
85%
12%
5%
88%
(f) EcoRI MspI (385−585bp)
47%
30%
13%
85%
(g) SphI MluCI (311−347bp)
46%
24%
2%
10%
(h) EcoRI SphI (280−342bp)
25%
19%
0.3%
2%
(i) PstI MseI (205−265bp)
65%
39%
0%
1%
(j) HindIII MspI (275−525bp)
19%
16%
0.2%
2%
(k) SbfI MspI (370−470bp)
21%
15%
0%
3%
(l) EcoRI MseI (335−535bp)
67%
21%
31%
38%
(m) EcoRI MspI (310−485bp)
27%
21%
0%
14%
(n) EcoRI MspI (273−347bp)
16%
13%
0%
0%
(o) EcoRI SphI (409−513bp
22%
17%
0%
0%
Proportion of fragments
Insert size

Supplementary Fig. 1: *Fragment size distribution predicted in in silico (black trend line) and observed in empirical data (red histogram). Black dashed line indicates two read lengths below which we expect overlapping reads. Black solid line indicates one read length below which we expect adaptor contamination. Grey box represents the intended insert size selection criteria chosen in the study. Percentages in black represent in silico prediction of the proportion of fragments and red are the proportion of fragments in empirical data.* ***(a)*** *Inbar et al (unpublished) did not use Bluepippin for size selection, only magnetic beads cleanup,* ***(b-c)*** *controlled experiments carried out on the same set of samples while switching the frequently cutting enzyme,* ***(d-o)*** *representative studies that used Bluepippin: (d) Baiz et al., 2018 (e) Trense et al., 2020 (f) Fritz et al., 2017 (g) Combs et al., 2017 (h) Termignoni et al., 2017 (i) Ivanov et al., 2018 (j) Jong et al., 2019 (k) Farleigh et al., 2020 (l) Ryan et al., 2017 (m) schley et al., 2020 (n) Portnoy et al., 2015 (o) Maigret et al., 2019*

Table 1: Table enumerating number of reads at each step of the pipeline for the controlled experiment.

| Enzyme | BfaI/EcoRI | MseI/EcoRI |
|---|---|---|
| # samples | 117 | 125 |
| Read pairs in raw data | 143,893,548 | 143,444,471 |
| Read pairs after removing extra samples from MseI (117 samples each) | 143,893,548 | 134,845,357 |
| Read pairs after removing reads from BfaI to equalise | 134,845,911 | 134,845,357 |

| | | | | |
|---|---|---|---|---|
| Adapter contamination based on multiqc results | 4.5% | | 20.2% | |
| # read pairs after trimmomatic | 120,712,950 | | 116,369,930 | |
| | Combined reads | Not combined read pairs | Combined reads | Not combined read pairs |
| # read pairs after Flash | 64,882,870 | 63,202,633 | 90,432,109 | 30,510,401 |
| % reads combined (overlaps) | 50.7% | | 74.8% | |
| # read pairs after Process_radtags | 64,958,995 | 61,432341 | 87,233,077 | 28,585,191 |

Table 2:

Enzyme pairs used in 66 studies that were examined in the literature survey

| Enzyme 1 | Enzyme 2 |
|---|---|
| EcoRI (28) | MspI (10), SphI (8), MseI (2), SbfI (2), Taqα1 (4), NlaIII (1), PstI (1) |
| SbfI (13) | MseI (6), MspI (6), Sau3AI (1) |
| SphI (6) | MluCI (5), MspI (1) |
| PstI (6) | MseI (1), MspI (1), NlaIII (1), AclI (1), HhaI (1), Csp6I (1), |
| HindIII (1) | MspI (1) |
| MfeI (1) | NlaIII (1) |
| MluCI (7) | NlaIII (6), MseI (1) |
| Xhol (1) | MseI (1) |
| NlaIII (1) | AciI (1) |
| Nsil (1) | MseI (1) |

Table 3: ddgRADer calculations

| User input | Calculation | output |
|---|---|---|
| Genome. choose enzyme pairs OR ask for enzyme recommendations | Fragment size distribution | 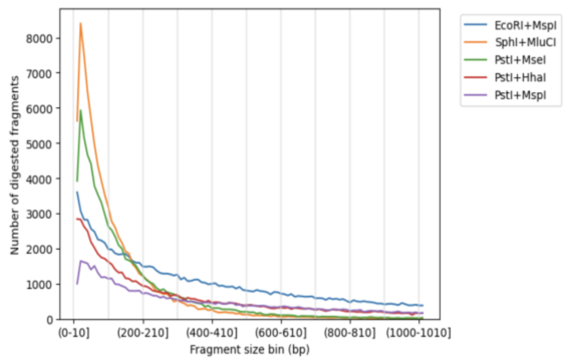 |
| For demographic analysis, enter desired # of SNPs; for genomic scan, enter desired SNP density | | |
| Size selection criteria | No. fragments x bin size*<br>* or double the read length; when bin size is > 2 x read length | No. basepairs that will be sequenced in the in silico digested genome |
| Expected SNP density | No. sequenced basepairs x expected SNP density | No. SNPs that will be genotyped |
| Sequencing yield + desired de | $\frac{sequencing\ yield}{total\ number\ of\ genomic\ fragments\ in\ the\ size\ selected\ range \times depth}$ | No. samples multiplexable |
| | $\frac{genomic\ fragments\ x\ bin\ size}{sequencing\ yield\ x\ read\ length}$ x 100 | Sequencing efficiency |

Table 4: Multiple linear regression for adaptor contamination with an interaction

| Variables | Coefficients | Std. error | t value | P value |
|---|---|---|---|---|
| (Intercept) | 0.0077016 | 0.0304884 | 0.253 | 0.8069 |
| Proportion adaptor contamination in silico | -0.0245421 | 0.0677269 | -0.362 | 0.7265 |
| Width of the size selection window | -0.0001226 | 0.0001406 | -0.872 | 0.4084 |
| Proportion adaptor contamination in silico: width of the size selection window | 0.0007978 | 0.0002737 | 2.915 | 0.0194 * |

Table 5: Multiple linear regression for read overlaps without adaptor contamination, additive model

| Variables | Coefficients | Std. error | t value | P value |
|---|---|---|---|---|
| (Intercept) | -0.1347334 | 0.0936718 | -1.438 | 0.210 |
| Proportion of read overlaps in silico excluding adaptor contamination | 0.8672884 | 0.4511788 | 1.922 | 0.113 |
| Width of the size selection window | 0.0001751 | 0.0002383 | 0.735 | 0.496 |

ddRAD-seq protocol

Modifications added by A. Brelsford, A. Mastretta-Yanes, J. Leuenberger, R. Sermier. Based on protocols from Parchman et al. 2012 Mol. Ecol. and Peterson et al. 2012 PLOS ONE.

**1. Restriction Digest:**

1. Prepare master mix I (see below, 3 mL prepared per sample), mix by vortexing, and centrifuge.

| Reagent | Number of samples<br>1x |
|---|---|
| 10x CutSmart | 0.9 |
| Water | 1.9 |
| MseI | 0.1 |
| EcoRI | 0.1 |

2. Place 6 mL of sample DNA in each well of a plate (DNA should ideally be at a minimum concentration of 10 ng/mL and a maximum concentration of 100 ng/mL, but lower concentrations may still work).
3. Add 3 mL of the combined master mix I to each well.
4. The total reaction volume should be 9 mL. Cover and seal the plate, vortex, centrifuge and incubate at 37°C for 8 hours on a thermal cycler with a heated lid. Inactivate the restriction enzymes at 65°C for 20 minutes.

**2. Adaptor Ligation**

1. Thaw MseI and EcoRI adaptors. These adaptors should already be annealed.

2. Prepare master mix II (see below, 1.6 mL prepared per sample), mix by vortexing, and centrifuge. As above, it is best to prepare an extra 20% (1.2x/sample).

| Reagent | Number of samples 1x |
|---|---|
| 10x CutSmart | 0.26 |
| 100 mM ATP | 0.116 |
| Water | 0.0565 |
| MseI Y adapter (dual indexing) 10 mM | 1.0 |
| T4 DNA Ligase | 0.1675 |

3. Add 1.6 mL to each well of the restriction digested DNA.
4. Add 1 mL of the EcoRI adaptor to each well (a unique barcoded adaptor for each DNA sample).
5. The total reaction volume should now be 11.6 mL. Cover and seal the plate, vortex, centrifuge and incubate at 16° C for 3 hours on a thermal cycler.
6. Dilute the Restriction-Ligation reaction with 38.4 mL of water

**3. Purification (short fragment removal) using Agencourt AMPure**

1. Let the AMPure bottle at room temperature for 30 min.
2. Shake the AMPure bottle gently to resuspend the magnetic beads. Vortex several minutes (1-2 min).
3. Add the correct volume of AMPure beads according to the ratio you want to use (we use a 0.8:1 ratio for this step) to the DNA volume.

4. Mix by pipetting 10 times or vortex 10 sec if it’s done in a tube.
5. Incubate 5 min at room temperature.
6. Place the reaction plate onto a magnet plate for 10 min to separate the beads from the solution. Be sure that the solution is clear before going to the next step.
7. Still on the magnet plate, aspirate the cleared solution (supernatant) and discard. Be careful not to move the plate or move/touch the beads.
8. Still on the magnet plate, dispense 200uL of 70% ethanol (freshly prepared same day from absolute ethanol) to each well and incubate at room temperature 1min (at least 30sec).
9. Still on the magnet plate, aspirate out the ethanol and discard
10. Still on the magnet plate, repeat steps 8 and 9.
11. Still on the magnet plate, let the plate at room temperature for 10-15 minutes to remove all traces of ethanol. Not too long; avoid over-drying the beads.
12. Remove the plate from the magnet plate. Add 40 mL of elution buffer (water, TRIS) by pipetting the mix 10-30 times.
13. Put the plate on the magnet plate again for 1 minute to separate the beads from the solution.
14. Still on the magnet plate, transfer the eluent with DNA into a new plate.

### 4. PCR Amplification

1. This PCR step uses the Illumina PCR primers to amplify fragments that have our adapters + barcodes ligated onto the ends. To ameliorate stochastic differences in PCR production of fragments in reactions, **we run four separate 10 mL** reactions per restriction-ligation product, and later combine them. If your sequencing batch includes fewer than 96 individuals, run each PCR at double volume (20 mL) to produce sufficient library quantity.

2. Prepare master mix III, vortex and centrifuge.

| Reagent | Number of samples 1x |
|---|---|
| Water | 2.15 |
| Q5 buffer | 2 |
| dNTP **(25mM)** | 0.08 |
| PCR Primer Mix | 0.67 |
| Q5 Taq | 0.1 |
| High GC enhancer | 2 |

3. Add 7 mL of the combined master mix III to each well of a plate.
4. Add 3 mL of the diluted restriction-ligation purified with AMPure mix.
5. Thermal cycler profile for this PCR: 98° C for 30s; 20 cycles of: 98° C for 20s, 60° C for 30s, 72° C for 40s; final extension at 72° C for 2 min.
6. Prepare master mix IV (see below, 1 mL per sample).

| Reagent | Number of samples 1x |
|---|---|
| Water | 0.05 |
| Q5 Buffer | 0.2 |
| PCR primer mix | 0.67 |
| dNTP **(25 mM)** | 0.08 |

7. Add 1 mL to each PCR product, run thermalcycler profile as follows: 98° C for 3 min, 60° C for 2 min, 72° C for 12 min.

**4. Repeat purification with Ampure beads**

**5. Size selection using Bluepippin**
